# Possible Magneto-Mechanical and Magneto-Thermal Mechanisms of Ion Channel Activation by Iron-Loaded Ferritin in Magnetogenetics

**DOI:** 10.1101/540351

**Authors:** Mladen Barbic

## Abstract

The palette of tools for stimulation and regulation of neural activity is continually expanding. One of the new methods being introduced is magnetogenetics, where mechano-sensitive and thermo-sensitive ion channels are genetically engineered to be closely coupled to the iron-storage protein ferritin. Such genetic constructs could provide a powerful new way of non-invasively activating ion channels in-vivo using external magnetic fields that easily penetrate biological tissue. Initial reports that introduced this new technology have sparked a vigorous debate on the plausibility of physical mechanisms of ion channel activation by means of external magnetic fields. I argue that the initial criticisms leveled against magnetogenetics as being physically implausible were possibly based on the overly simplistic and unnecessarily pessimistic assumptions about the magnetic spin configurations of iron in ferritin protein. Additionally, all the possible magnetic-field-based mechanisms of ion channel activation in magnetogenetics might not have been fully considered. I present and propose several new magneto-mechanical and magneto-thermal mechanisms of ion channel activation by iron-loaded ferritin protein that may elucidate and clarify some of the mysteries that presently challenge our understanding of the reported biological experiments. Finally, I present some additional puzzles that will require further theoretical and experimental investigation.

## Introduction

Interaction of biological systems with magnetic fields has puzzled and fascinated the scientific community for a long time^1–7^. While experimental evidence for magnetic sense in animals seems uncontroversial, the mystery of biophysical mechanism of its action remains unresolved. Despite the challenges in deciphering the fundamental operating principles of magnetic control of biological ion channels/cells/organisms, the attraction of influencing biological systems with magnetic fields has remained strong. This is mainly due to the fact that external DC and AC magnetic fields easily penetrate biological tissue, are easily generated by current carrying wires or permanent magnets, and their properties and engineering design tools are well understood. These features of magnetic fields are commonly used in medical diagnostics applications such as Magnetic Resonance Imaging (MRI)^8^, and there is a strong impetus to apply the same advantages of magnetic fields to control biological function, as is the case in Transcranial Magnetic Stimulation (TMS)^9^.

Coupling modern genetic engineering techniques with the magnetic fields to influence biological activity promises a particularly potent way to combine the strength of both methods in control of cellular function. This is the approach of a recent technique development, commonly termed magnetogenetics, where thermo-sensitive and mechano-sensitive ion channels are genetically engineered to be closely spatially coupled to the iron-storage protein ferritin^10–13^. Initial reports that introduced this new technology have received significant attention and commentary^14,15^, as well as considerable criticism^16^. The plausibility of presented mechanisms of mechanical and thermal ion channel has been challenged on physical grounds, and the present state of reported experimental observations in magnetogenetics and basic magnetic physics arguments that challenge those observations remain in conflict.

Here, I will present the case that the physical viability of magnetogenetics depends critically on many physical parameters of the basic control construct (iron-loaded ferritin protein coupled to the thermo-sensitive or mechano-sensitive ion channel in the cell membrane) that are presently not well known or understood, and therefore the physical possibility of magnetogenetics cannot yet be discounted and needs to be further explored. These critical parameters include the magnetism and magnetic spin configurations of iron atoms in the ferritin protein, as well as the realistic thermal, mechanical, and diamagnetic properties of ion channels and neural cell membranes coupled to the iron loaded ferritin. Additionally, I will argue that all the possible magnetic-field-based mechanisms of ion channel activation by iron-loaded ferritin in magnetogenetics might not have been fully considered. I will present and propose several new possible mechanisms of ion channel activation based on the magneto-caloric effect, mechanical cell membrane deformation by the diamagnetic force, and mechano-thermal Einstein-de-Haas effect. I will also discuss the fundamental magnetic moment fluctuations of the magnetic particle in ferritin and its presently unknown but potentially relevant effect on the ion channels in cell membranes.

## Magnetism of Iron in Ferritin Protein

The fundamental genetically engineered construct in magnetogenetics, as reported by the original articles^10–13^, is shown diagrammatically in Figure 1. Thermo-sensitive or mechano-sensitive ion channels of the TRP family of channels (TRPV1 and TRPV4) were genetically engineered to be closely spatially coupled to the novel chimeric iron-loading protein ferritin. External DC or AC magnetic fields were applied to this construct in in-vivo experimental settings, and it was preliminarily concluded that thermal or mechanical magnetic effects were likely responsible for the observed biological responses, as was intended by the genetic engineering methods. Meister has argued^16^ that the presently known magnetic force, torque, and heating mechanisms are many orders of magnitude smaller than necessary to activate ion channels and that the fundamental thermal energy, k_B_T, is much larger than the energy of any mechanism of magnetogenetics so far proposed. Meister’s criticism is fundamentally based on his assumptions on the number of iron atoms (N=2400) in the ferritin protein and on the magnetic configuration in ferritin protein that assumes iron spins as a collection of independent non-interacting particles (paramagnetic configuration). Here I will explore what the physical consequences are if those assumptions are expanded to include reasonable and experimentally observed larger number of iron atoms in the ferritin protein^17^ (N=4500) and spin configurations that allow for more strongly coupled iron spins within the ferritin core (ferromagnetic configuration).

**Figure 1.**
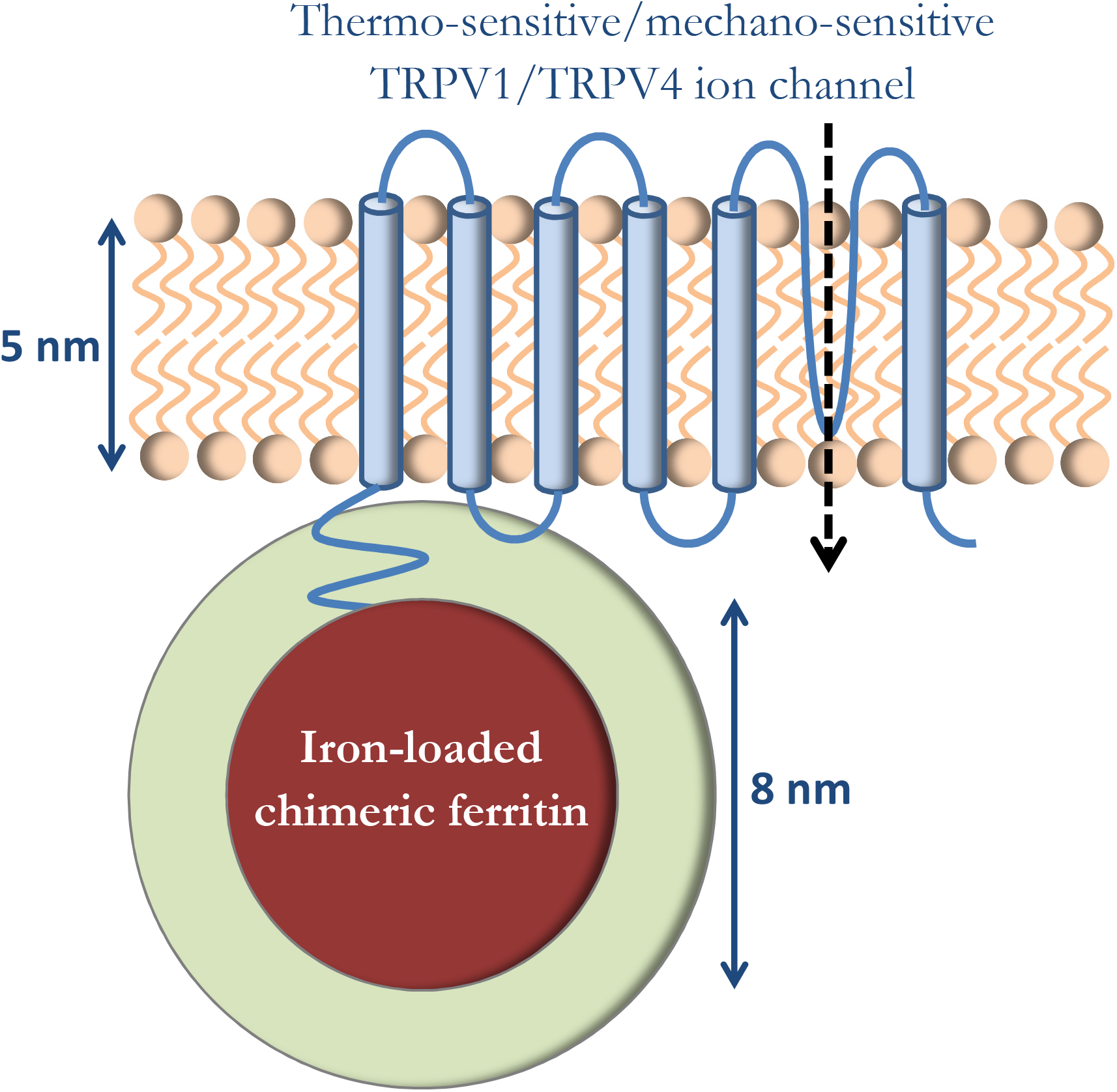
Genetically engineered construct in magnetogenetics, where thermo-sensitive or mechano-sensitive ion channel is closely coupled to the iron-loading protein ferritin. External DC or AC magnetic fields are applied to the construct in order to influence ion channel function.

It should first be recognized that magnetic properties of iron are notoriously structure sensitive^18^, and assigning the magnetic moment value to the iron atom between 0μ_B_ to 5μ_B_ (where μ_B_ is the Bohr magneton, μ_B_=9.27×10^-24^ Am^2^) and spin coupling configuration (paramagnetic, ferromagnetic, ferrimagnetic, antiferromagnetic, etc.) is highly variable and chemistry dependent. Along the same lines, it should be pointed out that the systematic and complete analysis of magnetism of the iron loaded ferritin core in all the reported magneto-genetics articles has not been performed, and therefore the magnetism of the fundamental magneto-genetic ferritin construct is not presently known. It is generally believed that iron forms different mineral forms within ferritin (for example, magnetite Fe_3_O_4_ or maghemite γFe_2_O_3_). The original enhanced iron-loading chimeric ferritin, reported by Iordanova et al.^19^, was characterized only by NMR and MRI, and showed that one of the fused heavy H and light L subunits constructs (the L*H chimera) exhibited significantly enhanced iron loading ability. It is not clear from the Iordanova et al. report how many iron atoms are indeed loaded into the L*H chimera construct, what chemical structure iron forms in that construct, what the spin configuration of the iron atoms in that construct is, or if there is any shape or crystalline or other magnetic anisotropy in the L*H ferritin (as the magnetic moment vs. magnetic field or magnetic moment vs. temperature measurements or chemical crystal analysis were not reported). In my analysis I will use the commonly stated maximum possible number of iron atoms in ferritin N=4500 and assume atomic moment of m_Fe_=5μ_B_ per iron atom (the highest value reported for iron in oxide from, Table 3.5 in Reference 18), with the understanding that the actual number of iron atoms in ferritin in all the magnetogenetics reports is not presently known.

Figure 2 diagrammatically shows the three distinct spin coupling configurations of iron atoms in ferritin that I will consider as reasonable possibilities. Figure 2a represents N irons spins in a conventional paramagnetic state where all the spins are magnetically independent from one another and non-interacting. This is the configuration considered by Meister^16^. Figure 2c shows the case of ferromagnetic coupling between N iron spin where all the spins are strongly coupled by the magnetic exchange interaction and magnetically behave as a single macro-spin in what is commonly termed a superparamagnetic state of the magnetic particle^18^. In Figure 2b I consider a state that lays in between these two extremes. In this configuration the N iron spins are separated into n independent clusters of N/n exchange coupled spins, and I will call this spin configuration a clusterparamagnetic state. For simplicity of analysis, I will assume that all of the n magnetic clusters in this spin configuration of the particle have the same number N/n of iron spins, and that the magnetic moments of the clusters within the ferritin particle are magnetically independent (non-interacting). It should be clear that Figure 2b represents the most general spin configuration with n clusters of N/n spins in each cluster, while the paramagnetic state of Figure 2a is a special case with n=N and the superparamagnetic state of Figure 2c is a special case where n=1. I will assume the physiological temperature of T=37°C=310K throughout. The thermal average magnetic moment in external magnetic field B of a single cluster of N/n iron spins in the ferritin particle is classically described by the Langevin function^18^:

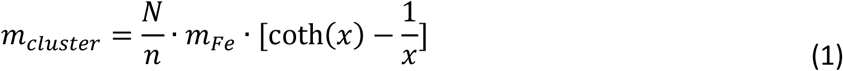

where parameter 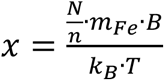, and the total thermal average magnetic moment m_TOT_ of the ferritin particle along the field direction is:

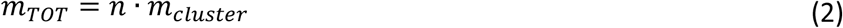

**Figure 2.**
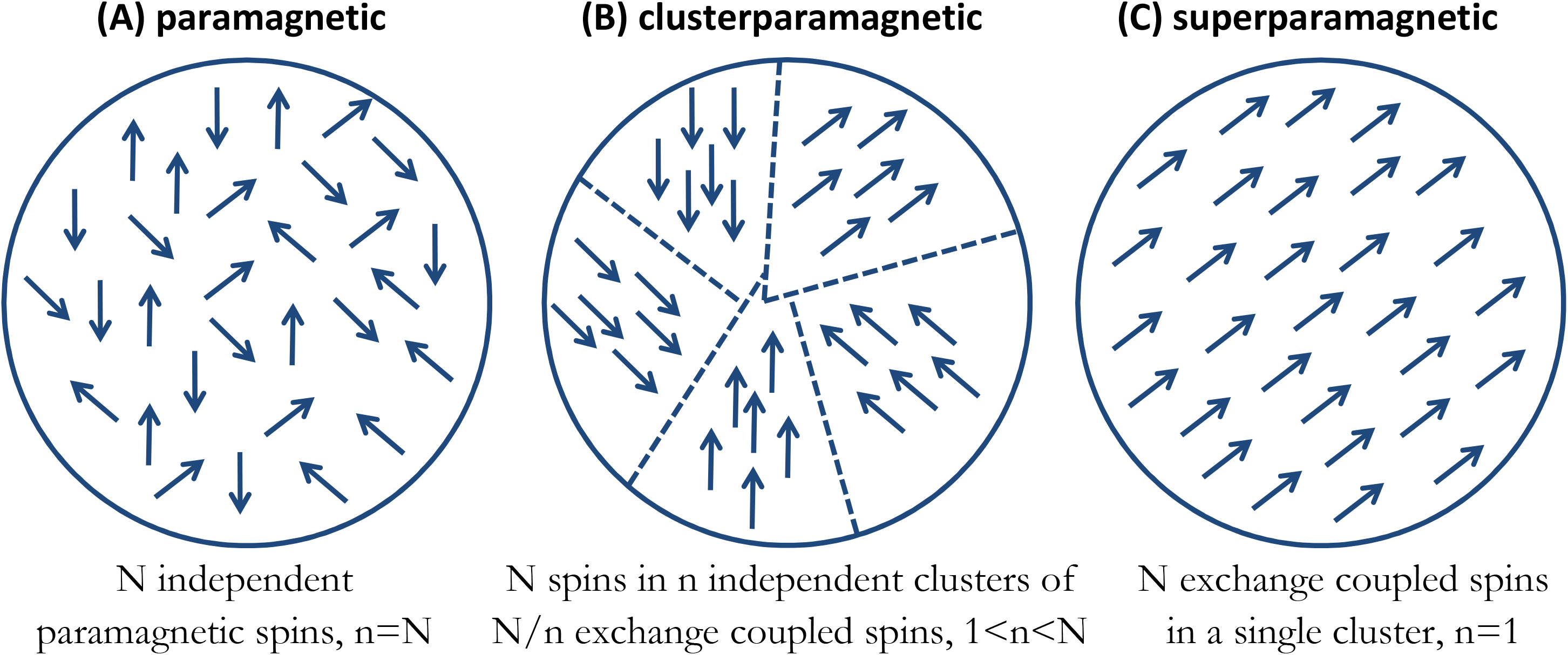
Three distinct spin coupling configurations of iron atoms in ferritin. (a) Paramagnetic state where N irons spins are magnetically independent from one another and non-interacting. (b) Clusterparamagnetic state where N iron spins are separated into n independent clusters of N/n exchange coupled spins. (c) Superparamagnetic state where all N spins are strongly coupled by the magnetic exchange interaction and behave as a single macro-spin.

Figure 3a shows the resulting ferritin particle magnetic moment vs. magnetic field at T=310K as a function of different cluster numbers and for the reasonable range of experimentally attainable laboratory magnetic fields (0 to 2 Tesla) generated by either electro-magnets, superconducting magnets, or permanent magnets.

**Figure 3.**
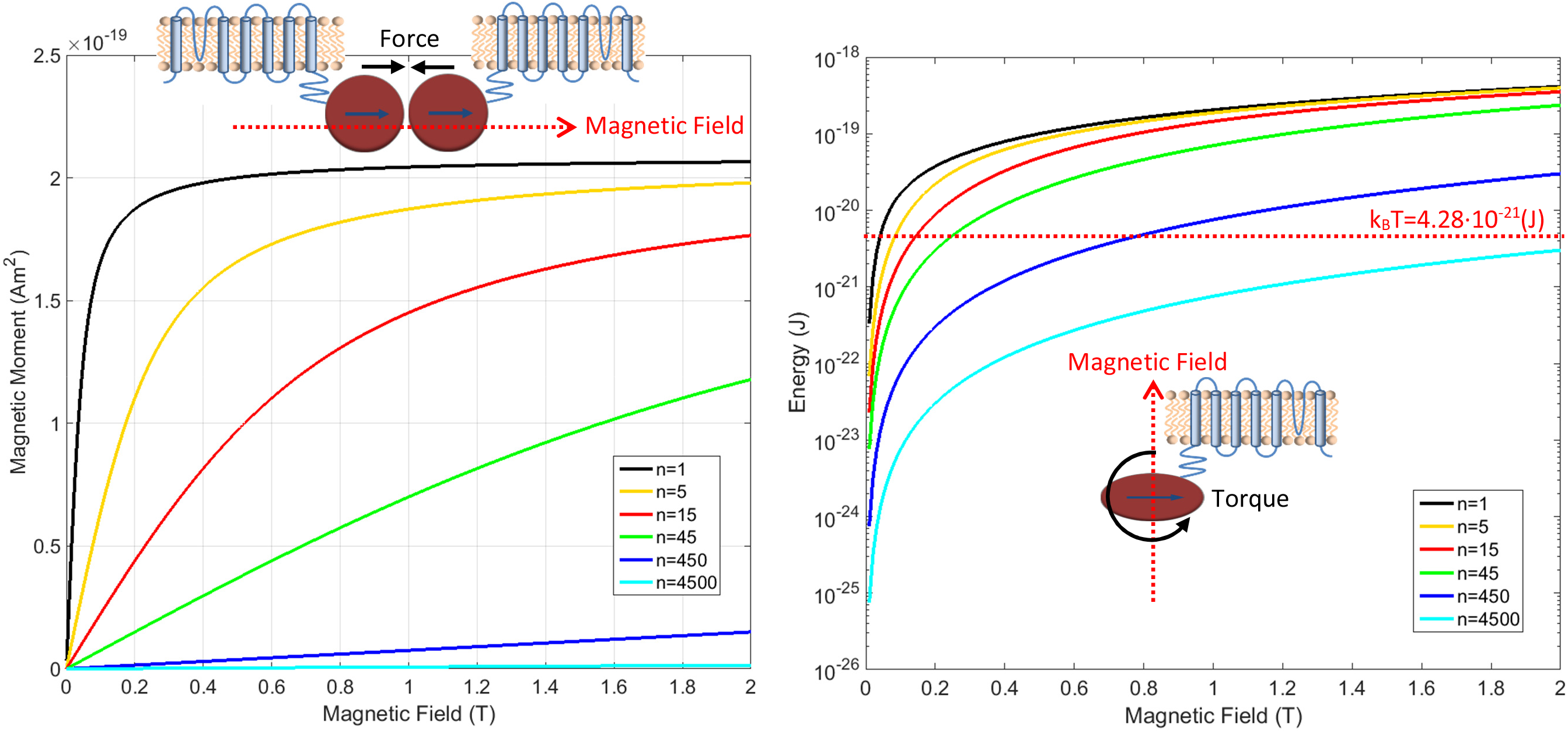
(a) Ferritin magnetic moment vs. magnetic field as a function of different cluster numbers n. For the superparamagnetic spin arrangement (n=1, black curve), the particle moment saturates at relatively low magnetic fields and the magnetic moment is three orders of magnitude higher than for the paramagnetic state (n=4500, light blue curve). The curves for the clusterparamagnetic configurations (n=5, 15, 45, 450) are also shown. The inset in (a) shows the attractive force configuration between two ferritins. (b) The interaction energy magnitude (E=m·B) of the iron loaded ferritin in the external magnetic field. For modest clustering of iron spins in ferritin the interaction energy is above k_B_T in moderate magnetic fields. The maximum theoretically possible torque on an anisotropic ferritin particle 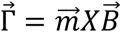, shown diagrammatically in the inset of (b), has interaction energy above k_B_T.

It is immediately apparent that the assumption about the spin configurations of iron atoms in the ferritin particle has dramatic effects on the total magnetism of the particle. For the paramagnetic (Figure 2a) spin arrangement of the particle (n=N=4500, light blue curve in Figure 3a) the magnetic susceptibility is low, and as an example, the total magnetic moment of the particle at a representative field of 1 Tesla is m=2.4×10^-22^ (Am^2^), in line with the numbers presented by Meister^16^. However, for the superparamagnetic (Figure 2c) spin arrangement of the particle (n=1, black curve of Figure 3a), the particle moment saturates at relatively low magnetic fields, well below 1 Tesla, where the magnetic moment has the value of m=2×10^-19^ (Am^2^), three orders of magnitude higher than for the paramagnetic state. This is the classic superparamagnetic particle behavior ^20^ where N=4500 spins are uniformly exchange coupled and act as a giant single paramagnetic spin. The other curves in Figure 3a show the magnetic moment for the clusterparamagnetic spin configurations (Figure 2b) of the particle (with cluster number n=5, 15, 45, 450) indicating that even modest clustering of N=4500 spins into n clusters of exchange coupled N/n spins can significantly increase the magnetic moment of the ferritin particle in reasonable laboratory magnetic fields.

While it is true, as Meister states^16^, that most experimental reports on ferritin magnetism indicate paramagnetic particles, that is not necessarily the case for the ferritin construct reported by Iordanova et al. and all the subsequent magnetogenetics reports for which the magnetization curves of the particles are not reported. In fact, there have been experimental reports on iron-loaded ferritin where the magnetization measurement closely follows the superparamagnetic curve (black line in Figure 3a). In the present analysis, the saturation magnetic moment of a superparamagnetic particle of 4500 iron atoms with moments of 5μ_B_ per atom is m_TOT_ = 22500μ_B_ = 2.1×10^-19^ (Am^2^) which is very much in line with the numbers in experiments reported by Bulte et al.^21^ and Moskowitz et al.^22^ indicating that the superparamagnetic state of iron-loaded ferritin is indeed possible. It is also instructive to consider the saturation magnetization M_s_ of magnetite in my calculation by dividing the particle saturation moment m_TOT_ by the volume V of the 8nm diameter particle (M_s_=m_TOT_/V) which results in M_s_=7.8×10^5^ (A/m). Commonly reported bulk magnetite magnetization is slightly lower at M_s_=4.8X10^5^ (A/m)^18^, but it should be noted that several independent experimental reports^23–25^ indicate that magnetite magnetization increases significantly in the size range below 10nm and reaches the value of M_s_=1X10^6^ (A/m). Therefore, my value of M_s_=7.8X10^5^ (A/m) for 8nm diameter ferritin particle of N=4500 iron spins with 5 μ_B_ per iron atom falls well within the experimentally reported range for magnetite.

## Force and Torque on Iron-Loaded Ferritin

It is instructive to reconsider the possible forces in magnetogenetics on a superparamagnetic ferritin particle (and therefore on the mechano-sensitive ion channel) and compare it to the paramagnetic ferritin particle case presented by Meister^16^. In the most optimistic case where the particle is for example at the entrance to the bore of a superconducting MRI magnet, where the magnetic field gradient could be on the order of ∇B=50 (T/m), the force on the paramagnetic ferritin would be on the order of *F* = *m* · ∇*B* =10^-20^(N), while the force on the superparamagnetic particle would be on the order of *F*=10^-17^ (N). The minimum required force to mechanically activate ion channels^26^ is on the order of 1pN=10^-12^ N. Therefore, the conclusion by Meister^16^ remains valid that the force from an externally applied magnetic field gradient on iron loaded ferritin, even the superparamagnetic one, is too low to be effective.

However, the situation is entirely different when one considers the mutual attractive force between two iron loaded ferritin particles that are both in a superparamagnetic spin configuration, as shown diagrammatically in the inset of Figure 3a. In a reasonable external magnetic field of 1 Tesla, both ferritin particles will be magnetically saturated, as Figure 3a shows. For 8nm diameter (radius r=4nm) ferritin particle, the maximum magnetic field gradient on the surface of the particle is:

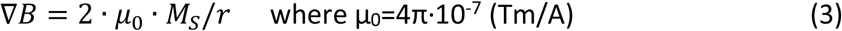

and has an enormous value of ∇B=4.9×10^8^ (T/m), much larger than anything achievable with external laboratory magnetic field gradients. Such a large magnetic field gradient from one ferritin particle acting on the second ferritin particle (and therefore on the mechanosensitive ion channels coupled to the ferritin particles) results in the maximum possible force on the order of 10^-10^(N)=100(pN), well above the required level for activating an ion channel. Therefore, if the construct reported by Iordanova et al. and used by magnetogenetics reports is similar in its magnetism or even more enhanced than what was reported by other superparamagnetic ferritin results of Bulte et al.^21^ and Moskowitz et al.^22^, then there is at least the theoretical plausibility that the two ferritin particles pulling on each other could result in a sufficient force to activate mechano-sensitive ion channels to which the ferritin particles are coupled.

Figure 3b shows the interaction energy magnitude (E=m·B) of the iron loaded ferritin configurations of Figure 2 in the external magnetic fields between 0-2 Tesla. On the same semilog plot I indicate the thermal energy level at the physiological temperature of T=310K, k_B_T=4.28×10^-21^ (J). As Meister has pointed out for the paramagnetic ferritin particle^16^ (n=4500 light blue curve in the plot) that interaction energy is lower than k_B_T. However, it is interesting that even for modest clustering of N=4500 iron spins into say n=450 clusters of N/n=10 exchange coupled spins each (n=450 dark blue curve in the plot) the interaction energy of the particle rises above k_B_T in a reasonable laboratory field of 1 Tesla. The magnetic energy diagram of Figure 3b is interesting in that the maximum theoretically possible torque on an anisotropic ferritin particle, as Meister has pointed out^16^, can easily be evaluated since torque 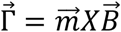. Therefore, there is in principle a theoretical possibility that an external magnetic field can exert a sufficient torque on an anisotropic ferritin particle (on the order of 10^-19^ N·m in a 1 Tesla external field) to activate mechano-sensitive ion channels.

## Diamagnetic Force on Ion Channel from Iron-Loaded Ferritin

In addition to the external forces on the iron loaded ferritin (and therefore on the coupled mechano-sensitive TRP ion channel) due to externally applied magnetic fields and field gradients, I now consider a case that to my knowledge has not been previously considered: the force due to the magnetic fields and field gradients from the ferritin particle itself on the intrinsically diamagnetic mechano-sensitive ion channel and neural cell membrane. I argue that this diamagnetic repulsive force could be sufficient to mechanically deform the ion channel and affect its function, as I diagrammatically describe in Figure 4.

**Figure 4.**
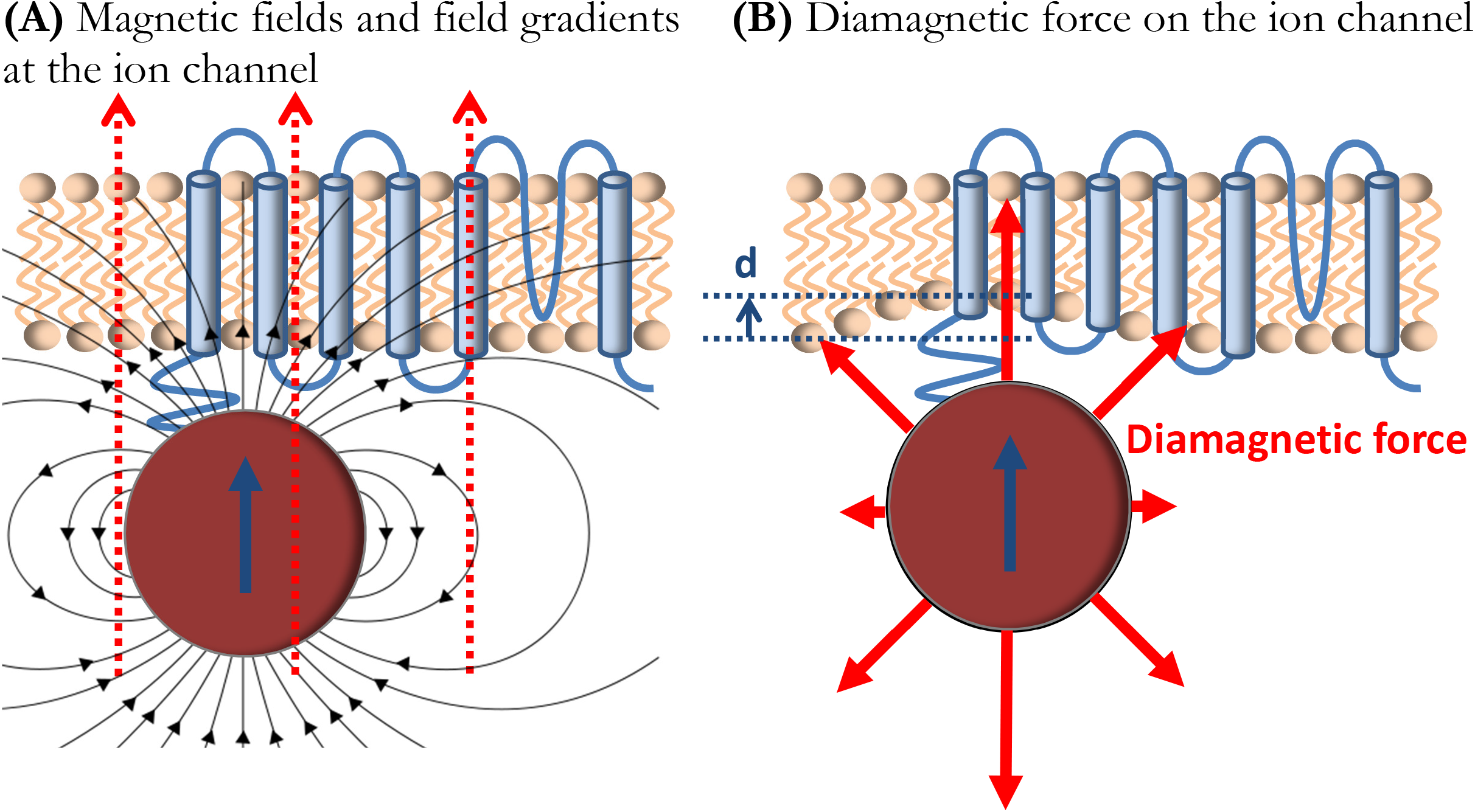
Diamagnetic force deformation of ion channel and cell membrane. (a) Diamagnetic ion channel in the cell membrane experiences magnetic fields from the ferritin particle and the externally applied magnetic field, as well as the large magnetic field gradient from the ferritin particle. This results in the repulsive diamagnetic force on the ion channel and the cell membrane in (b) that is sufficient to mechanically deform them and affect ion channel function.

Diamagnetism is generally considered the feeblest forms of known magnetism^18,27^, but it can have surprising and dramatic mechanical effects on biological materials (most of which are diamagnetic), such as levitation^28,29^ or restriction of water flow^30,31^, if the sufficient and necessary conditions of high magnetic fields and field gradients are present. The diamagnetic force per unit volume on a diamagnetic material is^29^:

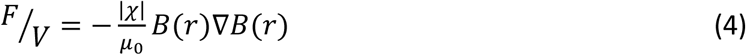

It is apparent that the critical parameter for generating appreciable force on a material with diamagnetic susceptibility χ is the product of the magnetic field and the magnetic field gradient *B*(*r*)∇*B*(*r*). In laboratory settings at the edge of a superconducting magnet bore this parameter can have values on the order of 10^3^ (T^2^/m)^29^.

I now consider what the value of this parameter is at the location of the ion channel/cell membrane that is closely coupled to the superparamagnetic ferritin particle, as shown in Figure 4a. In an externally applied uniform magnetic field of 1 Tesla the superparamagnetic particle magnetic moment will be saturated (as shown in Figure 3a), and the ion channel in cell membrane will experience the total combined magnetic field from the saturated magnetite particle B_particle_ and the external magnetic field B. The magnetic field on top of the saturated 8nm diameter magnetite particle in my model is:

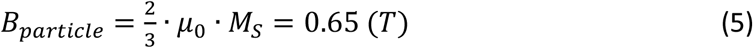

and the total maximum field seen by the ion channel and cell membrane is 1.65 (T). The ion channel in the cell membrane will also be under the influence of the particle gradient magnetic field of 4.9X10^8^ (T/m). This results in the critical parameter in the diamagnetic force calculations of *B∇B*=8.1×10^8^ (T^2^/m), many orders of magnitude larger than anything available in the common laboratory settings^29^. This combination of magnetic fields and magnetic field gradients from the ferritin particle will generate a repulsive diamagnetic force on the ion channel/cell membrane, as shown diagrammatically in Figure 4b. It is particularly intriguing that the diamagnetic repulsive force from the ferritin particle is exerted on the mechanosensitive ion channel in a neural cell membrane which is mechanically extremely soft^32^. The consensus that is emerging from the mechanical properties of ion channels and cell membranes experiments^33–35^ is that neural cells appear to have extremely low Young’s modulus, E, on the order of E=100 (Pa), the lowest of any known materials. The final parameter in the diamagnetic force equation (4) is the diamagnetic susceptibility χ of the ion channel and the cell membrane. It is reasonable to assume that this number is similar to that of water^18^, on the order of χ=-1×10^-5^, but one should not dismiss the possibility that ordered lipid cell membrane domains could have an order of magnitude higher value of diamagnetic susceptibility^36,37^.

Estimating the resulting deformation of the ion channel and cell membrane, as shown in Figure 4b, due to the diamagnetic force of Equation 4 is extremely difficult, as both the magnetic fields and the magnetic field gradients from the magnetite particle are extremely spatially varying in both direction and magnitude. However, preliminary 3D finite element modeling of the cell membrane deformation using the listed magnetic, mechanical, and diamagnetic parameters for the ferritin construct and the cell membrane in my model gives the estimate that the deformation d in Figure 4b is on the order of ~1nm (I thank Richard Smith for performing the numerical computer finite element modeling of this effect). Such deformation represents a significant volume fraction change of the overall cell membrane thickness of ~5nm, and is highly likely to modify the functional properties of mechano-sensitive ion channel in magnetogenetics.

## Magneto-Caloric Effect in Iron-Loaded Ferritin

In addition to the forces and torques that could possibly affect mechano-sensitive ion channels in magnetogenetics, heating of iron loaded ferritin by AC magnetic fields has also been considered as one of the mechanisms for thermo-sensitive ion channel activation and argued by Meister to also be implausible. I now consider a thermal mechanism that to my knowledge has also so far not been considered for the clusterparamagnetic spin arrangement in magnetogenetics: the DC magneto-caloric effect in the iron-loaded ferritin particle on the thermo-sensitive ion channel. Magneto-caloric effect refers to the heating and cooling of magnetic materials by DC magnetic fields^18^. It is fundamentally based on the physical principle that an ensemble of magnetic spins in zero magnetic field is fluctuating and therefore maximally randomized and in a high entropy state. Upon application of a polarizing magnetic field the magnetic spins align in the field which lowers the entropy of the spin ensemble. In an adiabatic process where there is no exchange of heat with the environment, this change in spin ensemble entropy has to be compensated for by the exchange of energy between the spin ensemble and the lattice, resulting in the change of temperature of the sample. This process of magnetic adiabatic cooling has been used in low-temperature physics for a long time^38^ to obtain milliKelvin temperatures in paramagnetic salt powders and sub-milliKelvin temperatures with nuclear spins in metals^39^. Although such magnetic adiabatic energy transfers are generally performed only at cryogenic temperatures, McMichael et. al.^40^ have pointed out that this process can be potentially performed at higher temperatures if the spins are coupled into superparamagnetic clusters (such as I diagrammatically show in Figure 2b). In fact, these authors have shown that, for a given magnetic field B and temperature T, there is an optimum clusterparamagnetic size that will generate maximum entropy change and energy transfer from the spin ensemble to the lattice. This hypothesis seems to have been experimentally confirmed^41,42^. Here I investigate the magneto-caloric energy transfer values for the specific case of clusterparamagnetic iron loaded ferritin (Figure 2) with N=4500 iron atoms with 5μ_B_ atomic moments at a physiological temperature of T=310K and reasonable laboratory magnetic fields.

Figure 5 diagrammatically shows the basic process. In zero magnetic field, n clusterparamagnetic moments (configuration of Figure 2b with N/n spins in each cluster) of the particle are randomly fluctuating (Figure 5a) and have a high magnetic entropy state. Upon application of an external magnetic field on the order of 1T, the clusterparamagnetic moments will mostly align with the magnetic field (Figure 5b) and go into a low entropy state. McMichael et al.^40^ have derived the classical entropy change in such as process for n-cluster particle (with N/n spins in each cluster):

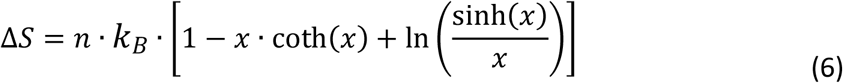

where, again, parameter 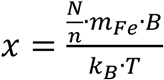

**Figure 5.**
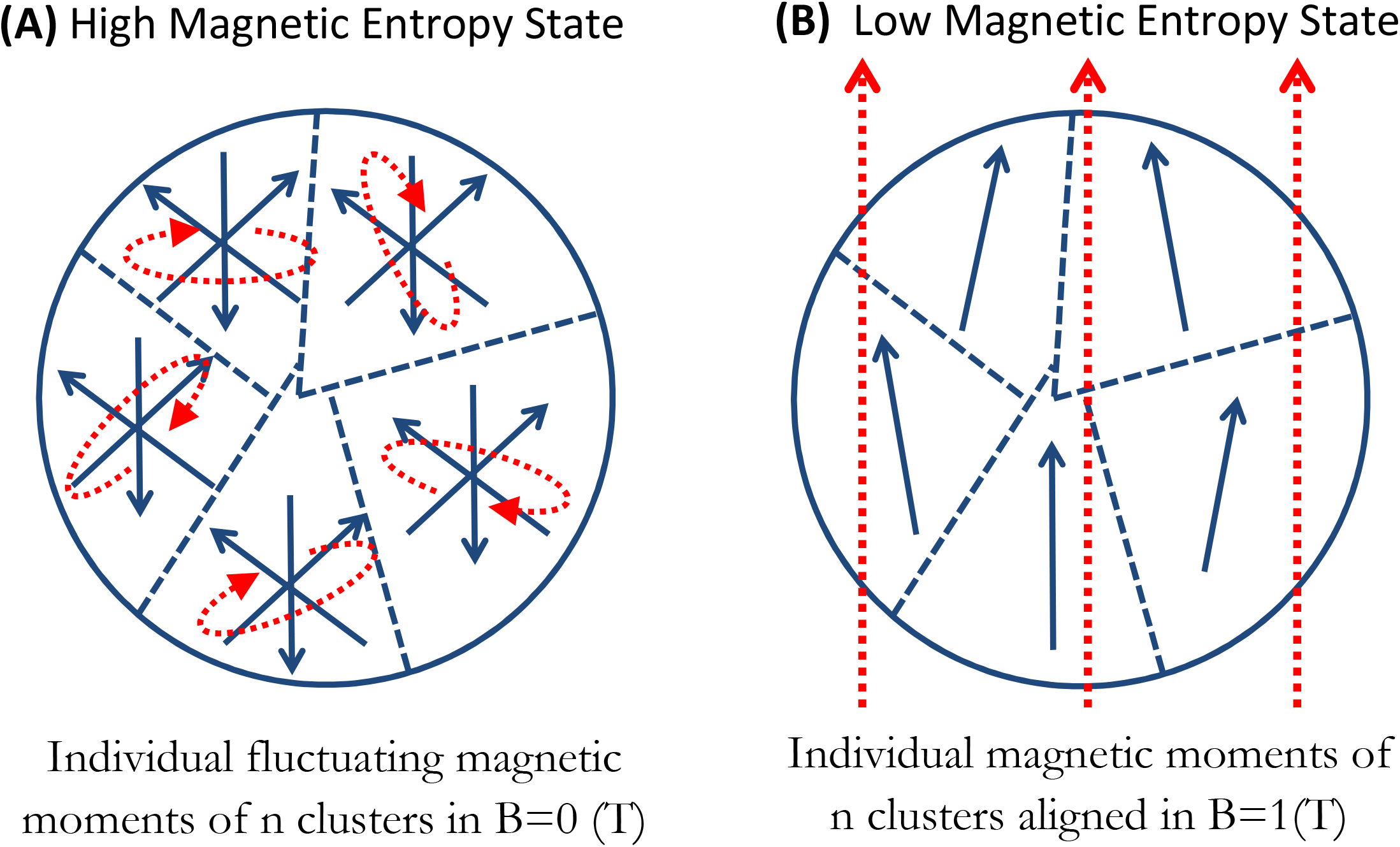
Magneto-caloric effect in clusterparamagnetic ferritin. (a) In zero magnetic field, clusterparamagnetic moments are randomly fluctuating and have a high magnetic entropy state. Upon application of external magnetic field in (b), the clusterparamagnetic moments align with the field and have low entropy. In the adiabatic process this change in spin entropy is compensated for by the exchange of energy between the spin ensemble and the lattice. For a given magnetic field B and temperature T, there is an optimum clusterparamagnetic size that w0ill generate maximum entropy change and energy transfer.

This magnetic entropy change results in the energy transfer to the lattice of Δ*E* = *T* · Δ*S*. Figure 6 shows the numerical calculation results for that magneto-caloric energy change in N=4500 iron atom ferritin particle as a function of the number of clusters n in the particle at physiological temperature of T=310K and several reasonable magnetic field values. As McMichael et al.^40^ have shown, for a given magnetic field B and temperature T there is an optimum clusterparamagnetic size that will generate maximum entropy change and energy transfer to the lattice. For example, for physiological temperature of 310K and magnetic field change of 1 Tesla (black points in Figure 5), the ferritin particle with N=4500 iron spins that separate into n=15 equal clusters of N/n=300 exchange coupled spins in each cluster will provide the maximum energy transfer to the lattice. In the same plot I show the thermal energy k_B_T level that reveals the interesting features in this analysis. For the paramagnetic spin state of the ferritin particle (N=n=4500) no value of magnetic field is sufficient to achieve magneto-caloric particle energy change near the level of k_B_T. However, if the ferritin particle of N=4500 iron spins clusters into n=100 (or fewer) clusters of N/n exchange coupled spins each, the magneto-caloric energy transfer from the spin ensemble to the lattice at physiological temperatures and reasonable laboratory magnetic fields is higher than the k_B_T level, including for the superparamagnetic state (Figure 2c, n=1 point in Figure 6). It is not clear how this magneto-caloric energy transfer from the clusterparamagnetic spin ensemble to the thermosensitive or mechano-sensitive ion channel could occur for channel activation, but the energy scales in the magneto-caloric process indicate that it is theoretically feasible.

**Figure 6.**
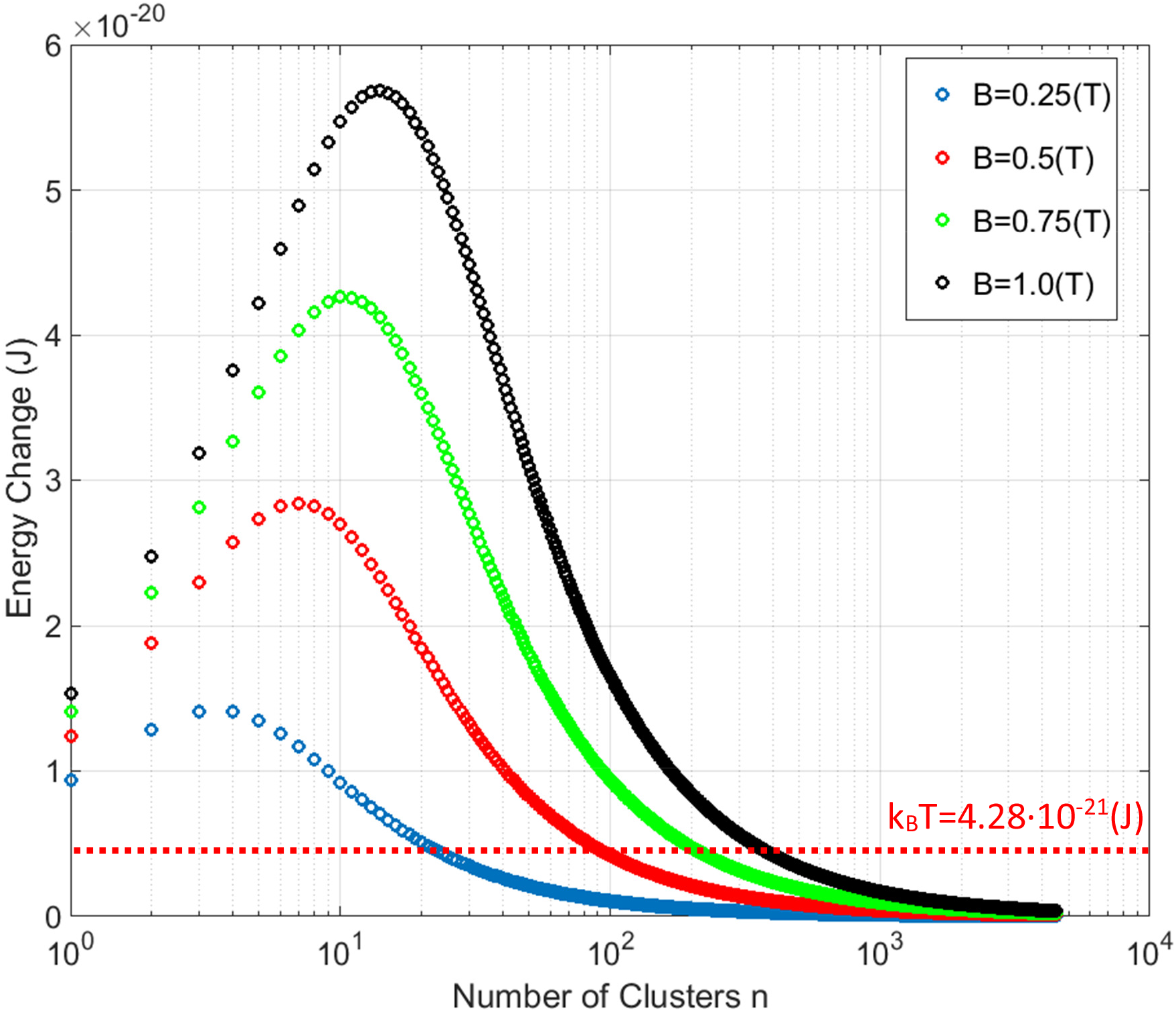
Magneto-caloric energy change for N=4500 iron atom ferritin particle as a function of the number of clusters n at a physiological temperature T=310K and several applied magnetic field values. For a given applied magnetic field B and temperature T there is an optimum clusterparamagnetic size that will generate maximum entropy change and energy transfer to the lattice. No value of applied magnetic field is sufficient to achieve magneto-caloric energy change above k_B_T for a paramagnetic ferritin (n=4500). However, grouping of the spins into exchange coupled clusters achieves magneto-caloric energy changes above the k_B_T level.

It might well be, as pointed out by Meister^16^, that this amount of energy transfer localized to the single ferritin particle is not sufficient to locally heat up the thermo-sensitive ion channel. However, it may also be that, as discussed by Keblinski et al.^43^, a large number of iron-loaded ferritin particles expressed throughout the neural cell membrane generate a global temperature rise (on a longer timescale) that is orders of magnitude larger than the negligible short timescale temperature rise adjacent to a single iron-loaded ferritin protein.

## Einstein-de Haas Effect on Iron-Loaded Ferritin

I now present another potential magneto-mechanical mechanism, the Einstein-de Haas effect ^44,45^, that appears feasible in iron-loaded ferritin and that to my knowledge has not been previously considered for possible ion channel activation. It is a fundamental tenet of quantum mechanics that magnetic moment m of a particle is proportional to mechanical angular momentum L of that particle: m=γL where γ is the gyromagnetic ratio γ=e/m_e_ for spin angular momentum of iron (where e is charge of the electron and me is mass of the electron^18^). The Einstein-de Haas effect refers to the magneto-mechanical effect, required by the conservation of angular momentum, where a reversal of a magnetic moment of a sample by an applied magnetic field has to be accompanied by a corresponding change in mechanical angular momentum of that sample (interesting historical perspective on the Einstein-de Haas effect is given in Chapter 2 of Reference 45). This physical principle has been experimentally confirmed on both the macroscopic^46^ and microscopic samples^47^, and recently on the molecular scale as well^48^. Here, I numerically evaluate the mechanical and thermal consequences of the Einstein-de Haas effect on the iron-loaded ferritin protein and therefore on the mechano-sensitive and thermo-sensitive ion channels in magnetogenetics.

Figure 7 shows diagrammatically the basic Einstein-de Haas principle. In the initial state magnetic field is applied along the positive z-axis and the magnetic moment +m is aligned with the magnetic field. This moment fundamentally caries mechanical angular momentum of L=+m/γ. As the magnetic field is reversed, the magnetic moment also reverses to −m value, which corresponds to the new mechanical angular momentum of L=-m/γ. This total change of angular momentum of ΔL=2m/γ has to, by the law of conservation of angular momentum, be compensated for by the mechanical rotation of the magnetite particle^49^ (the Einstein-de Haas effect). This change in mechanical angular momentum is proportional to the change in rotational kinetic energy^49^ of the magnetite particle 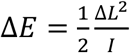, where I is the rotational moment of inertia of a spherical magnetite particle of radius 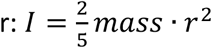. Assuming the density of magnetite of 5.24×10^3^ (kg/m^3^), the mass of the 8nm diameter particle is mass=1.4×10^-21^(kg), and the energy transferred to the magnetite particle by the rotational motion imparted by the magnetic moment reversal is Δ*E*=0.33×10^-21^(J). This rotational kinetic energy of the magnetite particle due to the Einstein-de Haas effect has to eventually transfer to the lattice through friction.

**Figure 7.**
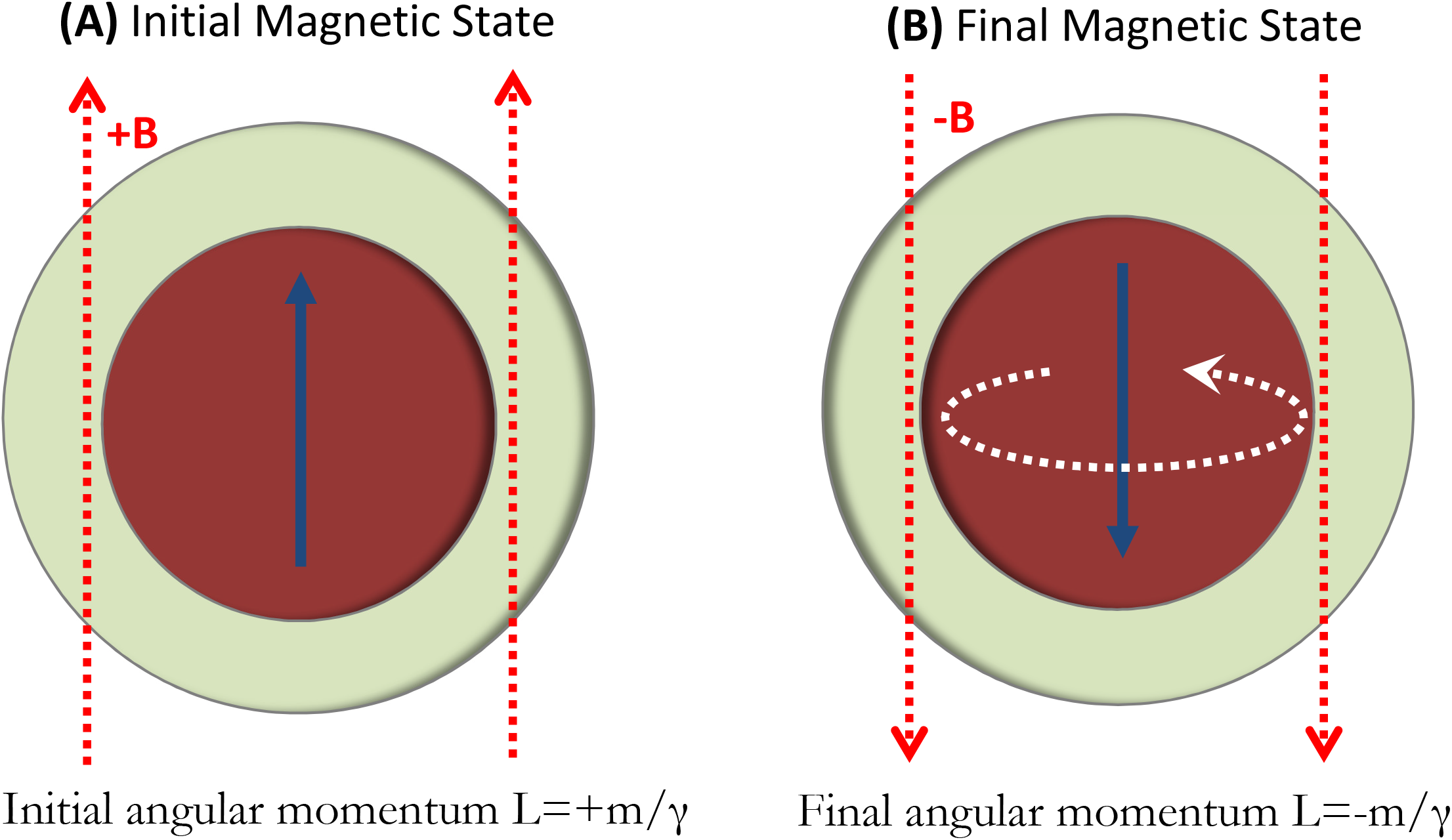
Einstein-de Haas effect in ferritin. (a) Ferritin magnetic moment +m is aligned with the field and caries mechanical angular momentum of L=+m/γ. Magnetic moment reversal to −m results in the total change of angular momentum of ΔL=2m/γ that is compensated for by the mechanical rotation of the particle or by the mechanical torque on the particle.

It is important to note that the resulting rotational energy of 0.33×10^-21^ (J) by a single magnetite particle moment reversal (one half of the applied AC magnetic field cycle) is a significant fraction of k_B_T=4.28×10^-21^ (J). It is interesting to compare this value to the maximum energy loss per cycle of 1 (J/kg) typically reported for magnetic hyperthermia applications^50,51^. For the Einstein-de Haas effect presented here, this value for a full single cycle is 2·Δ*E*/mass=0.47(J/kg). Therefore, it would appear that the Einstein-de Haas effect should be considered on par with the typical Brown and Néel relaxation modes in magnetic particle hyperthermia^52–54^ in contributing to the sample heating. As was the case for the magneto-caloric effect, it is not yet known how this Einstein-de Haas magneto-mechanical process would transfer energy to the thermo-sensitive and mechano-sensitive ion channel for activation, but the energy scales in this process are again very close to k_B_T.

This entire analysis of the Einstein-de Haas magneto-mechanical frictional heating for a magnetite particle inside the ferritin protein was predicated on the assumption that the magnetite particle is free to move inside the protein cage, a condition that to my knowledge is not presently known. So one should also consider the situation in which the spherical magnetite particle is fixed inside the ferritin protein and cannot freely rotate. In this situation the change in magnetic moment direction of the magnetite particle from +m to −m which results in the change of the mechanical angular momentum L of the particle from +m/γ to −m/γ now has to impart a torque on the surrounding medium 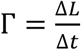, where Δ*t* is the time of reversal of the magnetite particle magnetic moment. A reasonable value to assume for the time of the moment reversal is Δ*t* =1(nsec), which results in the torque of 2m/(γ·A*t*)=2.4×10^-21^ (N·m) per half cycle of the applied AC magnetic field. The energy scale of this torque is again in the same range as k_B_T=4.28×10^-21^ (J). How this Einstein-de Haas torque would be transferred to the mechano-sensitive ion channel for activation is unknown and remains to be theoretically and experimentally explored.

## Magnetic Moment Fluctuations of Iron-Loaded Ferritin

I finally discuss a topic that also appears not to have been previously considered, which is the effect of superparamagnetic ferritin particle magnetic moment fluctuations and its potential effect on the ion channels. As discussed earlier, for a superparamagnetic spin arrangement (Figure 2c) of the ferritin particle, the magnetic field near the particle surface (Figure 4a) is relatively large at 0.65 Tesla. This field from the particle is not static, but is in fact fluctuating rapidly in time^55,56^, as I show schematically in Figure 8a. The frequency of this fluctuation at physiological temperature is significant and estimated to be in the GHz frequency range. Such superparamagnetic moment fluctuations have been experimentally observed^57^, even on a single particle scale at low temperatures^58–60^. Therefore, the ion channel in the vicinity of the iron loaded ferritin experiences large magnetic field gradients and Tesla-scale magnetic fields at GHz-scale frequencies, as well as the corresponding GHz frequency diamagnetic forces and torques, as discussed earlier. Upon application of the external DC magnetic field, of say 1 Tesla, the saturation of the magnetic moment in that external DC field results in the ion channel experiencing a Tesla-scale DC magnetic field, diamagnetic force, and torque as I show in Figure 8b. I am presently not aware of any experimental or theoretical studies that have investigated cell membrane or ion channel behavior in such extreme conditions of combined high amplitude magnetic fields (Tesla-scale) and field gradients (10^8^-10^9^ T^2^/m) and ultra-high frequencies (1-10 GHz scale), and yet that would appear to be the environment in which the ion channels in magnetogenetics constructs are operating. Therefore, studying ion channel properties in those conditions, both theoretically and experimentally, would be warranted before final conclusions could be made about the possibilities and limitations of magnetogenetics.

**Figure 8.**
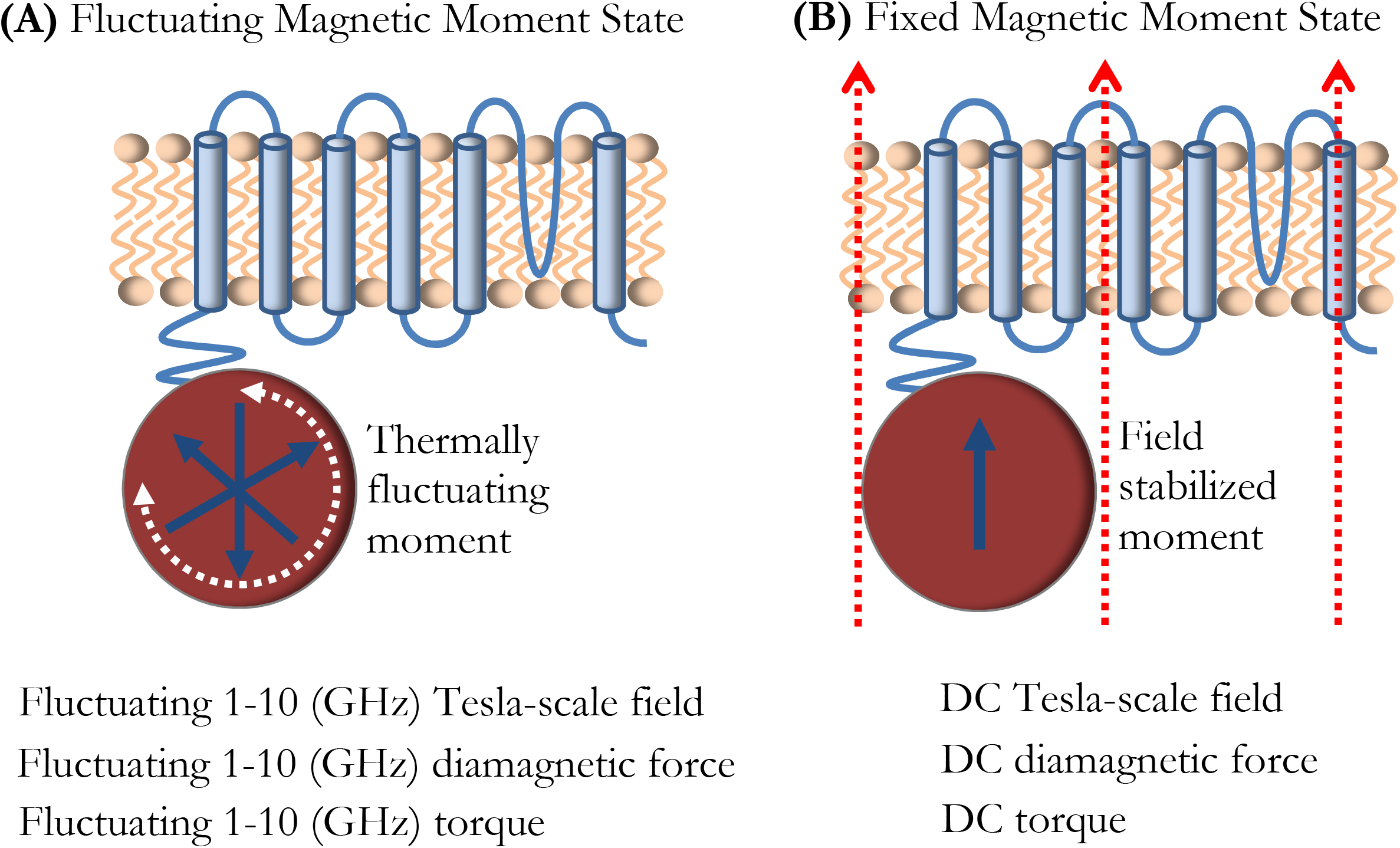
In zero external field, the ion channel experiences Tesla-scale magnetic fields and large field gradients from the fluctuating superparamagnetic ferritin moment at GHz-scale frequencies, as well as the corresponding AC diamagnetic forces and torques. In the external field, the ion channel experiences Tesla-scale DC magnetic fields and large field gradients from the stabilized ferritin magnetic moment, and the corresponding DC diamagnetic forces and torques.

## Conclusion

I have presented several physical mechanisms of magneto-thermal and magneto-mechanical interactions in the settings relevant to iron-loaded ferritin particle-based magnetogenetics that to my knowledge have previously not been considered. I have found that several of these interactions have energy scales on the order of or higher than the k_B_T level, and that the energies associated with these interactions depend strongly on the iron spin arrangement assumptions within the ferritin particle. Many parameters that are critical to the full understanding of all the prospects and viabilities of magnetogenetics remain unknown and include: (1) the exact iron magnetism, spin configuration and magnetic anisotropy of the ferritin construct utilized so far in magnetogenetics studies, (2) the realistic mechanical and thermal nano-environment around the iron-loaded ferritin, (3) the diamagnetic (and perhaps even paramagnetic) properties of ion channels and cell membranes next to the iron-loaded ferritin, and (4) the functional properties of ion channels under the influence of simultaneously large amplitude and ultra-high frequency magnetic fields and field gradients. It is entirely possible that additional magnetic effects on ion channels in neural cell membranes, such as the phase changes and deformations in lipid bilayers due to magnetic fields^27,61–63^, gradient magnetic field effects on the ion diffusion^64^ and the resting membrane potential of cells^65^, as well as the induction of calcium influx in cells by nanomagnetic forces^66^ could further influence magnetogenetic activation of cells. As the technical advances in genetically-encoded bio-mineralization of ferritin continue^67,68^, further theoretical and experimental investigations of all of the possible mechanisms, parameters, and conditions in magnetogenetics should be pursued before the final verdict on the possibilities and limitations of this new neuro-stimulation technology is rendered.

## Acknowledgement

This work was supported by the Howard Hughes Medical Institute. I thank Tim Harris for allowing me to explore this research topic and Richard Smith for multiple discussions and preliminary finite element computer simulations of the diamagnetic force deformations of cell membranes. I acknowledge discussions on this topic with Jeffrey Friedman, Sarah Stanley, Leah Kelly, Tim Harris, Markus Meister, Mikhail Shapiro, Joe Kirschvink, Arnd Pralle, Scott Sternson, Loren Looger, Karel Svoboda, Jacob Robinson, Ali Güler, Manoj Patel, Alan Koretsky, Doug Morris, Stephen Dodd, and David Hunt. I emphasize that my acknowledgment of the discussions with these scientists does not necessarily equal their endorsement of the ideas and results I presented here on this controversial topic.

## References

1 Kirschvink, J. L. & Gould, J. L. Biogenic Magnetite as a Basis for Magnetic-Field Detection in Animals. Biosystems 13, 181–201 (1981).

2 Kirschvink, J. L., Walker, M. M. & Diebel, C. E. Magnetite-based magnetoreception. Curr Opin Neurobiol 11, 462–467 (2001).

3 Walker, M. M., Dennis, T. E. & Kirschvink, J. L. The magnetic sense and its use in long-distance navigation by animals. Curr Opin Neurobiol 12, 735–744 (2002).

4 Bazylinski, D. A. & Frankel, R. B. Magnetosome formation in prokaryotes. Nat Rev Microbiol 2, 217–230 (2004).

5 Johnsen, S. & Lohmann, K. J. The physics and neurobiology of magnetoreception. Nature Reviews Neuroscience 6, 703–712 (2005).

6 Johnsen, S. & Lohmann, K. J. Magnetoreception in animals. Phys Today 61, 29–35 (2008).

7 Hore, P. J. & Mouritsen, H. The Radical-Pair Mechanism of Magnetoreception. Annu Rev Biophys 45, 299–344, doi:10.1146/annurev-biophys-032116-094545 (2016).

8 McRobbie, D. W., Moore, E. A. & Graves, M. J. MRI from picture to proton. Third edition. edn, (University Printing House, Cambridge University Press, 2017).

9 Walsh, V. & Cowey, A. Transcranial magnetic stimulation and cognitive neuroscience. Nat Rev Neurosci 1, 73–79, doi:10.1038/35036239 (2000).

10 Stanley, S. A. et al. Radio-wave heating of iron oxide nanoparticles can regulate plasma glucose in mice. Science 336, 604–608, doi:10.1126/science.1216753 (2012).

11 Stanley, S. A., Sauer, J., Kane, R. S., Dordick, J. S. & Friedman, J. M. Remote regulation of glucose homeostasis in mice using genetically encoded nanoparticles. Nat Med 21, 92–98, doi:10.1038/nm.3730 (2015).

12 Stanley, S. A. et al. Bidirectional electromagnetic control of the hypothalamus regulates feeding and metabolism. Nature 531, 647–650, doi:10.1038/nature17183 (2016).

13 Wheeler, M. A. et al. Genetically targeted magnetic control of the nervous system. Nat Neurosci 19, 756–761, doi:10.1038/nn.4265 (2016).

14 Anikeeva, P. & Jasanoff, A. Problems on the back of an envelope. Elife 5, doi:10.7554/eLife.19569 (2016).

15 Nimpf, S. & Keays, D. A. Is magnetogenetics the new optogenetics? EMBO J 36, 1643–1646, doi:10.15252/embj.201797177 (2017).

16 Meister, M. Physical limits to magnetogenetics. Elife 5, doi:10.7554/eLife.17210 (2016).

17 Chasteen, N. D. & Harrison, P. M. Mineralization in ferritin: an efficient means of iron storage. J Struct Biol 126, 182–194, doi:10.1006/jsbi.1999.4118 (1999).

18 Coey, J. M. D. Magnetism and magnetic materials. (Cambridge University Press, 2010).

19 Iordanova, B., Robison, C. S. & Ahrens, E. T. Design and characterization of a chimeric ferritin with enhanced iron loading and transverse NMR relaxation rate. J Biol Inorg Chem 15, 957–965, doi:10.1007/s00775-010-0657-7 (2010).

20 Bean, C. P. & Jacobs, I. S. Magnetic Granulometry and Super-Paramagnetism. J Appl Phys 27, 1448–1452 (1956).

21 Bulte, J. W. et al. Magnetoferritin: characterization of a novel superparamagnetic MR contrast agent. J Magn Reson Imaging 4, 497–505 (1994).

22 Moskowitz, B. M. et al. Determination of the preexponential frequency factor for superparamagnetic maghemite particles in magnetoferritin. J Geophys Res-Sol Ea 102, 22671–22680 (1997).

23 Arora, S. K. et al. Giant magnetic moment in epitaxial Fe(3)O(4) thin films on MgO(100). Phys Rev B 77 (2008).

24 Orna, J. et al. Origin of the giant magnetic moment in epitaxial Fe3O4 thin films. Phys Rev B 81 (2010).

25 Guan, X. F., Zhou, G. W., Xue, W. H., Quan, Z. Y. & Xu, X. H. The investigation of giant magnetic moment in ultrathin Fe3O4 films. Apl Mater 4 (2016).

26 Milo, R. & Phillips, R. Cell biology by the numbers. (Garland Science, Taylor & Francis Group, 2016).

27 Yamaguchi, M. & Tanimoto, Y. Magneto-science: magnetic field effects on materials: fundamentals and applications. (Kodansha; Springer, 2006).

28 Geim, A. K., Simon, M. D., Boamfa, M. I. & Heflinger, L. O. Magnet levitation at your fingertips. Nature 400, 323–324 (1999).

29 Simon, M. D. & Geim, A. K. Diamagnetic levitation: Flying frogs and floating magnets (invited). J Appl Phys 87, 6200–6204 (2000).

30 Ueno, S. & Iwasaka, M. Properties of Diamagnetic Fluid in High-Gradient Magnetic-Fields. J Appl Phys 75, 7177–7179 (1994).

31 Ueno, S. & Iwasaka, M. Parting of Water by Magnetic-Fields. Ieee TMagn 30, 4698–4700 (1994).

32 Tyler, W. J. The mechanobiology of brain function. Nat Rev Neurosci 13, 867–878, doi:10.1038/nrn3383 (2012).

33 Lu, Y. B. et al. Viscoelastic properties of individual glial cells and neurons in the CNS. Proc Natl Acad Sci U S A 103, 17759–17764, doi:10.1073/pnas.0606150103 (2006).

34 Sanchez, D. et al. Noncontact measurement of the local mechanical properties of living cells using pressure applied via a pipette. Biophys J 95, 3017–3027, doi:10.1529/biophysj.108.129551 (2008).

35 Wu, P. H. et al. A comparison of methods to assess cell mechanical properties. Nat Methods 15, 491–498, doi:10.1038/s41592-018-0015-1 (2018).

36 Braganza, L. F., Blott, B. H., Coe, T. J. & Melville, D. The Superdiamagnetic Effect of Magnetic-Fields on One and 2 Component Multilamellar Liposomes. Biochim Biophys Acta 801, 66–75 (1984).

37 Ginzburg, V. L., Gorbatsevich, A. A., Kopayev, Y. V. & Volkov, B. A. On the Problem of Superdiamagnetism. Solid State Commun 50, 339–343 (1984).

38 Giauque, W. F. A thermodynamic treatment of certain magnetic effects. A proposed method of producing temperatures considerably below, 1(0) absolute. J Am Chem Soc 49, 1864–1870 (1927).

39 Simon, F. E. Low temperature physics. (Pergamon Press, 1952).

40 Mcmichael, R. D., Shull, R. D., Swartzendruber, L. J., Bennett, L. H. & Watson, R. E. Magnetocaloric Effect in Superparamagnets. J Magn Magn Mater 111, 29–33 (1992).

41 Mcmichael, R. D., Ritter, J. J. & Shull, R. D. Enhanced Magnetocaloric Effect in Gd3ga5-Xfexo12. J Appl Phys 73, 6946–6948 (1993).

42 Shao, Y. Z., Zhang, J. X., Lai, J. K. L. & Shek, C. H. Magnetic entropy in nanocomposite binary gadolinium alloys. J Appl Phys 80, 76–80 (1996).

43 Keblinski, P., Cahill, D. G., Bodapati, A., Sullivan, C. R. & Taton, T. A. Limits of localized heating by electromagnetically excited nanoparticles. J Appl Phys 100 (2006).

44 Einstein, A. Experimental detection of Amper’s molecular currents. Naturwissenschaften 3, 237–238 (1915).

45 Galison, P. How experiments end. (University of Chicago Press, 1987).

46 Scott, G. G. Review of Gyromagnetic Ratio Experiments. Rev Mod Phys 34, 102–& (1962).

47 Wallis, T. M., Moreland, J. & Kabos, P. Einstein-de Haas effect in a NiFe film deposited on a microcantilever. Appl Phys Lett 89 (2006).

48 Ganzhorn, M., Klyatskaya, S., Ruben, M. & Wernsdorfer, W. Quantum Einstein-de Haas effect. Nat Commun 7, 11443, doi:10.1038/ncomms11443 (2016).

49 Chudnovsky, E. M. Conservation of Angular-Momentum in the Problem of Tunneling of the Magnetic-Moment. Phys Rev Lett 72, 3433–3436 (1994).

50 Hergt, R. et al. Physical limits of hyperthermia using magnetite fine particles. Ieee T Magn 34, 3745–3754 (1998).

51 Hergt, R., Dutz, S., Muller, R. & Zeisberger, M. Magnetic particle hyperthermia: nanoparticle magnetism and materials development for cancer therapy. J Phys-Condens Mat 18, S2919–S2934 (2006).

52 Rosensweig, R. E. Heating magnetic fluid with alternating magnetic field. J Magn Magn Mater 252, 370–374 (2002).

53 Dutz, S. & Hergt, R. Magnetic nanoparticle heating and heat transfer on a microscale: Basic principles, realities and physical limitations of hyperthermia for tumour therapy. Int J Hyperthermia 29, 790–800, doi:10.3109/02656736.2013.822993 (2013).

54 Deatsch, A. E. & Evans, B. A. Heating efficiency in magnetic nanoparticle hyperthermia. J Magn Magn Mater 354, 163–172 (2014).

55 Brown, W. F. Thermal Fluctuations of a Single-Domain Particle. Phys Rev 130, 1677–+ (1963).

56 Brown, W. F. Thermal Fluctuations of Fine Ferromagnetic Particles. Ieee T Magn 15, 1196–1208 (1979).

57 Casalta, H. et al. Direct measurement of superparamagnetic fluctuations in monodomain Fe particles via neutron spin-echo spectroscopy. Phys Rev Lett 82, 1301–1304 (1999).

58 Wernsdorfer, W. et al. Experimental evidence of the Neel-Brown model of magnetization reversal. Phys Rev Lett 78, 1791–1794 (1997).

59 Piotrowski, S. K., Matty, M. F. & Majetich, S. A. Magnetic Fluctuations in Individual Superparamagnetic Particles. Ieee TMagn 50 (2014).

60 Hevroni, A., Tsukerman, B. & Markovich, G. Probing magnetization dynamics in individual magnetite nanocrystals using magnetoresistive scanning tunneling microscopy. Phys Rev B 92 (2015).

61 Maret, G. & Dransfeld, K. Macromolecules and Membranes in High Magnetic-Fields. Physica B & C 86, 1077–1083 (1977).

62 Tenforde, T. S. & Liburdy, R. P. Magnetic Deformation of Phospholipid-Bilayers – Effects on Liposome Shape and Solute Permeability at Prephase Transition-Temperatures. J Theor Biol 133, 385–396 (1988).

63 Kurashima, H., Abe, H. & Ozeki, S. Magnetic-field-induced deformation of lipid membranes. Mol Phys 100, 1445–1450 (2002).

64 Kinouchi, Y. et al. Effects of Static Magnetic-Fields on Diffusion in Solutions. Bioelectromagnetics 9, 159–166 (1988).

65 Zablotskii, V., Polyakova, T., Lunov, O. & Dejneka, A. How a High-Gradient Magnetic Field Could Affect Cell Life. Sci Rep 6, 37407, doi:10.1038/srep37407 (2016).

66 Tay, A., Kunze, A., Murray, C. & Di Carlo, D. Induction of Calcium Influx in Cortical Neural Networks by Nanomagnetic Forces. Acs Nano 10, 2331–2341 (2016).

67 Matsumoto, Y., Chen, R., Anikeeva, P. & Jasanoff, A. Engineering intracellular biomineralization and biosensing by a magnetic protein. Nat Commun 6, 8721, doi:10.1038/ncomms9721 (2015).

68 Liu, X. et al. Engineering Genetically-Encoded Mineralization and Magnetism via Directed Evolution. Sci Rep 6, 38019, doi:10.1038/srep38019 (2016).

